# Comparison of different extraction kits to isolate microRNA from *Galleria mellonella* (wax moth) larvae infected with *Metarhizium brunneum* (ARSEF4556)

**DOI:** 10.1101/606004

**Authors:** Muneefah A. Alenezi, Tariq M. Butt, Daniel C. Eastwood

## Abstract

MicroRNAs (miRNAs) play an important role in regulating gene expression and are involved in developmental processes in animals, plants and fungi. To understand the role of miRNAs in a biological system, it is important to optimise the extraction procedures to obtain high quality and quantity nucleic acid that enable high throughput sequencing and expression analysis. Numerous kit-based miRNA extraction protocols have been optimised generally to single cell or tissue cultures. Fungi, however, often occupy physically and chemically complex environments which miRNA make extraction challenging, such as fungal pathogens interacting within plant or animal host tissue. We used a *Galleria mellonella* (wax moth) larvae and entomopathogenic fungus *Metarhizium brunneum ARSEF 4556* host/pathogen model to compare commercially available miRNA extraction kits (Invitrogen PureLink™ miRNA Isolation Kit, Ambion mirVana™miRNA Isolation Kit and Norgen microRNA purification Kit). Our results showed reproducible and significant differences in miRNAs extraction between the kits, with the Invitrogen PureLink™ miRNA Isolation protocol demonstrating the best performance in terms of miRNA quantity, quality and integrity isolated from fungus-infected insect tissue.

## Introduction

Small RNA (sRNA) molecules have been increasingly recognised as significant factors regulating gene expression [1]. MicroRNAs (miRNAs) are an endogenous, 22-nucleotide, noncoding, single stranded RNA species that form a group of gene regulators involved in developmental processes in animals, plants and fungi [1, 2]. Ensuring the isolation of good quality miRNA samples is essential for downstream analysis, i.e. high throughput sequencing, with challenges associated with sample handling and miRNA extraction needing to be addressed [3]. Errors during sample handling (such as accidental contamination during the extraction process) and poor storage conditions can compound RNA quality-loss [4, 5]. As total RNA and miRNA are extracted in the same way, degraded total RNA will mean low miRNA concentration in a sample [6, 7]. Furthermore, low concentration of total RNA in a sample makes the estimation of miRNA abundance particularly difficult [8].

Extraction of miRNAs from samples can be technically challenging because of their small size and their attachment to cellular lipids and proteins [9–11]. Earlier studies on relatively low complex samples (e.g. single cell lines) have identified differences in quantity and quality of miRNA extracted with different commercial kits, with some highlighting the need for protocol optimisation [12, 13]. The success of commercial miRNA extraction kits on more complex systems consisting of a range of tissue types and/or multiple organisms are not well described, particularly comparing between treatments where samples change and deteriorate over time, e.g. host-pathogen interactions. In order to obtain miRNA from fungal pathogen, both the host tissue and fungal cells need to be homogenised and disrupted to release the nucleic acids.

*Metarhizium brunneum* ARSEF 4556 (previous name *M. anisopliae*) is a broad host range entomopathogenic fungus used as a biocontrol agent that undergoes morphogenic and physiological change during the infection process [14, 15]. A reproducible extraction protocol is required to investigate the potential regulation by miRNAs during pathogenesis. We tested three commonly used miRNA extraction protocols using a complex mixed system of *M. brunneum* against the insect host *Galleria mellonella* using both healthy and infected host tissue. *G. mellonella* is increasingly used as a model system to test microbial pathogenesis [16–19]. In this study, the interaction of *G. mellonella* larvae with *M. brunneum* provides a general fungal pathogen system with which to assess molecular protocols aimed at assessing fungi differentiating within living tissues.

The three protocols tested were PureLink™ miRNA Isolation Kit (Invitrogen), mirVana™miRNA Isolation Kit (Ambion) and microRNA purification Kit (Norgen). To the best our knowledge these kits have not been previously compared, and not for complex samples. We report that the quantity and quality of miRNA extracted varied significantly between the different extraction protocols. While extraction quality between *G. mellonella* healthy and *M. brunneum*-invaded tissue remained constant for any given protocol, the Invitrogen PureLink™ provided the greatest miRNA yield and quality from our samples.

## METHODS

### Fungal culture

*M. brunneum* (ARSEF 4556) was obtained from the Swansea University culture collection and grown on Sabourand dextrose agar (SDA, 40 gL^−1^ D-glucose, 10 gL^−1^ mycological peptone, 5 gL^−1^ technical agar (Sigma,UK), 0.5 gL^−1^ chloramphenicol) at 28 °C in the dark for 14 days to obtain the conidia. The conidia were harvested by using sterile distilled water containing 0.03% v/v Tween 80 and the concentration determined using a haemocytometer. Conidial viability was determined over a 122 hr time course using a plate count technique on SDA [20].

### Preparation and Inoculation of G. mellonella

*G. mellonella* (*Lepidoptera*) were maintained at 28°C in an artificial nutrition medium (15% (v/w) bee honey, 15% (w/w) wax, 15% (w/w) glycerol, 15% (w/w) fat free dry milk, and 40% (w/w) corn and wheat flour. Four *G. mellonella* larvae at 5-6^th^ stage [21] were submerged in 40 ml *M. brunneum* conidia suspension (1×10^8^ conidia ml^−1^) for 35 seconds, placed into Petri plates with moist filter paper and then sealed with Parafilm^®^ and incubated at 28°C. Control larvae were dipped into 0.03% (v/v) Tween 80 for 35 seconds and all treatments were repeated in triplicate. After incubation the larvae were frozen under liquid nitrogen and stored at −80 °C.

### MicroRNA and RNA Extraction

Invitrogen PureLink™ miRNA Isolation Kit, Ambion mirVana™miRNA Isolation Kit and Norgen microRNA kits were used to isolate miRNA and total RNA from *G. mellonella* larvae 72 hr post-infection with *M. brunneum*, uninfected *G. melonella* larvae and *M. brunneum* SDA-grown conidia (see Table 1 for kit overviews). Samples were prepared following manufacturer’s guidelines. Tissue, 100 mg, was used for Invitrogen PureLink™ miRNA Isolation and Ambion mirVana™miRNA Isolation kits (the whole *G. mellonella* larvae were used for both kits) and 50 mg tissue was used (half larva was used) for the Norgen microRNA purification kit. All samples were ground with a micropestle under liquid nitrogen and the standard protocol (frozen tissue extraction) was followed for each kit. The RNA was eluted in 100 μl RNase-free water for the Invitrogen PureLink™ miRNA Isolation kit, 100 μl elution buffer for the Ambion mirVana™ miRNA Isolation and 50 μl for the Norgen microRNA purification kit, and stored at −80°C.

**Table 1.** Overview of the miRNA and RNA isolation kits used in this study.

|  | **Invitrogen** | **Ambion** | **Norgen** |
| --- | --- | --- | --- |
|  | **PureLink** | **mirVana** | **MicroRNA** |
| **Molecules isolated** | Total RNA<br>inc small RNA<sup>#</sup> | Total RNA<br>inc small RNA | Total RNA<br>inc small RNA |
| **Quantity of biomass**<br>**required** | 100 mg | 100 mg | 50 mg |
| **Isolation chemistry** | Guanidine isothiocyanate<br>Ethanol precipitation | Phenol:Chloroform<br>Ethanol precipitation | Guanidine salt<br>Ethanol precipitation |
| **Column details** | Silica-based membrane,<br>2 columns (total & smRNA<br>extraction) | Glass fibre filter,<br>3 columns (total & smRNA<br>extraction) | Resin-based<br>membrane,<br>2 columns (total & smRNA<br>extraction) |
| **Cost per sample\*** | 3.36 GBP | 12.75 GBP | 10.8 GBP |
| **Steps in protocol** | 6 | 7 | 6 |
| **Protocol time per**<br>**sample** | 15 minutes | 30 minutes | 30 minutes |

<sup>#</sup> Small RNA = microRNA, small interfering RNA, tRNA and 5S RNA.
\* Purchased 2016 in Pounds Sterling (GBP) inc. taxes.

### RNA analysis

miRNAs and total RNA were quantified and integrity analyzed with the Agilent 2100 Bioanalyzer (Agilent Technologies, Santa Clara, CA) using the Agilent Small RNA chip and RNA pico-chip kits respecitvely. RNA concentration and purity was also measured at 260nm and 280nm absorbance using the Nanodrop spectrophotometer (NanoDrop Technologies, Wilmington, DE, USA). Data processing and analysis were conducted using GraphPad prism V5.0d software to compare the quantity and quality of microRNA. Molecular data sets were analyzed using two-way Analysis of Variance (ANOVA) with Tukey HSD post-test. Statistical analysis of the data was carried out in SPSS [22].

## Results

Invitrogen, Ambion and Norgen miRNA extraction kits were successfully used to isolate total RNA including miRNA from infected *G. mellonella* larvae with *M. brunneum* (72 hrs post-inculation), uninfected larvae (72 hrs) and *M. brunneum* culture. The quantity of miRNA isolated from the most complex sample (infected *G. mellonella*) showed significantly greater yield obtained from the Invitrogen PureLink kit (146.9 ng/μl, +/−5.1) measured by the Agilent Bioanalyzer compared with the Norgen MicroRNA (2.29 ng/μl, +/−0.434) or Ambion mirVana (0.773 ng/μl, +/−0.159) kits (Table 2). Similar results were obtained from uninfected *G. mellonella* and *M. brunneum* pure culture (Figure 1). In addition, the 260:280 and 260/230 absorbance ratios obtained using the Nanodrop showed that miRNA A260:A280 purity obtained with Invitrogen kit was better at 1.97 than that obtained with Norgen (1.77) or Ambion (1.51) kits, and A260:A230 values of 0.94 Norgen, 0.89 Ambion and 1.97 Invitrogen.

**Figure. 1.**
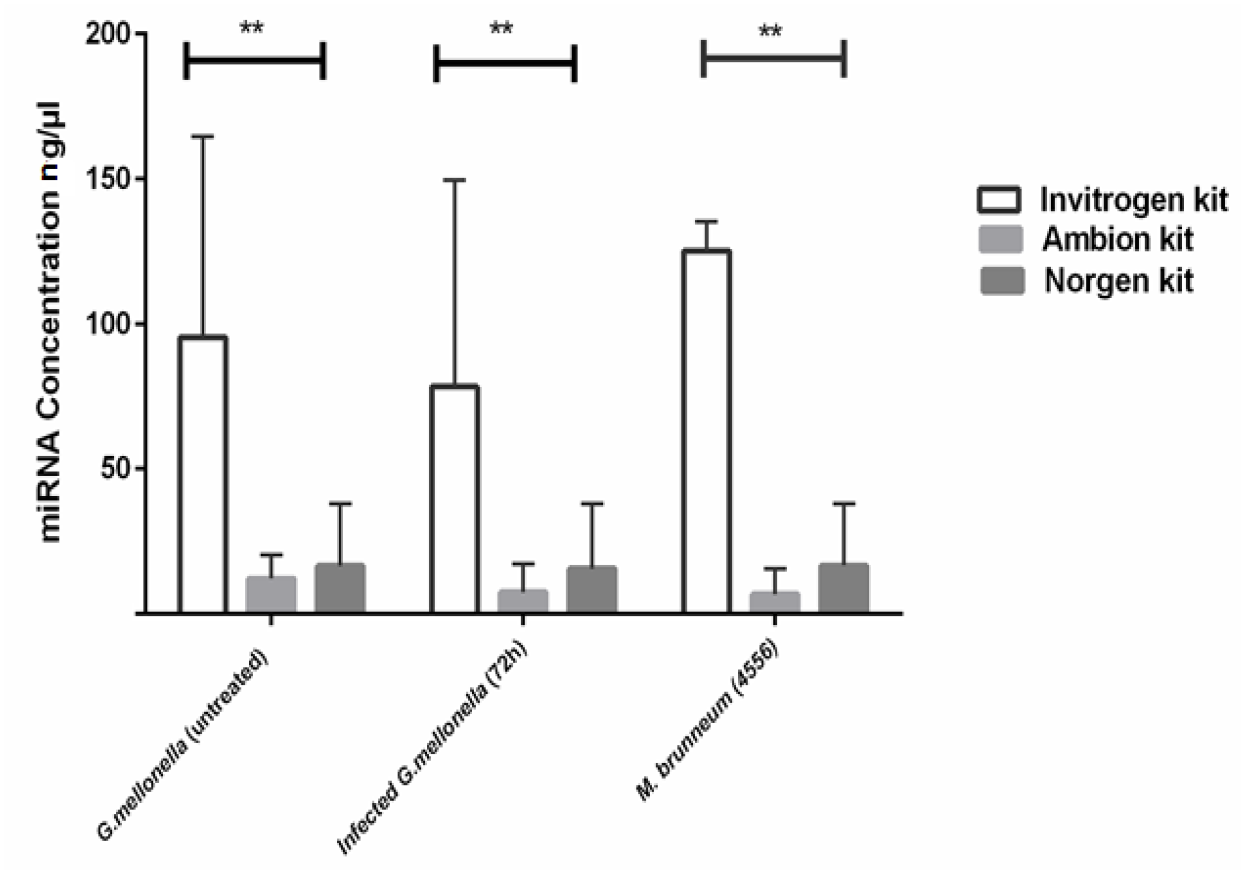
miRNA concentration extracted from *G. mellonella* and *M. brunneum* samples using the Aglient Bioanalyzer. The miRNA concentrations represent mean of three repeat extractions using the Invitrogen PureLink, Ambion mirVana, Norgen MicroRNA purification kits. Error bars shows significant differences (**) in the miRNA concentration (two way ANOVA, p<0.01) between samples processed by each of the kits.

**Table 2.** Assessment of small RNA extraction quality obtained from Invitrogen PureLink, Ambion mirVana and Norgen MicroRNA extraction kits from a complex sample of two interacting species: *Galleria mellonella* infected with *M. brunneum* for 72 hr were prepared in triplicate and values are presented as mean SD (range).

|  | Invitrogen kit | Ambion kit | Norgen kit |
| --- | --- | --- | --- |
| Small RNA concentration [ng/μl] | 117.1 (+/-18.2) | 1.6 (+/-0.0036) | 8.0 (+/-3.37) |
| miRNA concentration [ng/μl] | 146.9 (+/- 5.1) | 0.773 (+/-0.159) | 2.29 (+/-0.434) |
| miRNA range (ng/μl) | (141.8-151.2) | (0.574-0.892) | (1.856-2.724) |
| miRNA:small RNA ratio | 80:1 | 48:1 | 29:1 |

The quality of RNA obtained with the Invitrogen isolation kit (as indicated by the miRNAs and small RNA ratio of 80:1 using Bioanalyzer) was significantly higher than the 48:1 and 29:1 obtained by the Ambion kit and the Norgen kit respectively (Figure 2, ANOVA, p<0.005). Greater quality of miRNA was also obtained from the Invitrogen kit across all samples used, i.e. infected / uninfected *G. mellonella* samples and *M. brunneum* cultures. The miRNA fraction with sizes of approximately 18 nt and 30 nt measured by Bioanalyzer were of a higher purity for the Invitrogen kit than the other kits (Figure 3A). The Invitrogen kit appeared to yield miRNA with greater integrity when comparing the sizes of miRNA from the Bioanalyser-derived electropherograms (Figure 3A, B, and C), suggesting that the greater miRNA quanitiy obtained was in part due to lower degradation of the sample. The miRNA obtained using the Intvitrogen extraction protocol met the criteria (the percentage of miRNA in small RNA to assess the RNA quality) for further processing and high thoughput Illumina sequencing of the miRNAs present. The performance of the Ambion and Norgen kits appeared similar to one another with regards to the integrity of the miRNA obtained, i.e. evidence of degraded RNA in most samples (low RNA yield would result in failure of detection of miRNA present in low abundance). Figure 4 provides a representative example of the gel images obtained for samples extracted via each of the kits and shows the quality of miRNA verification. The relative quantity and quality of miRNA obtained from the more complex sample of *G. mellonella* larvae infected with *M. brunneum* for 72 hrs was comparable to that obtained from the non-infected larvae and *M. brunneum* culture controls (Figure 4C).

**Figure. 2.**
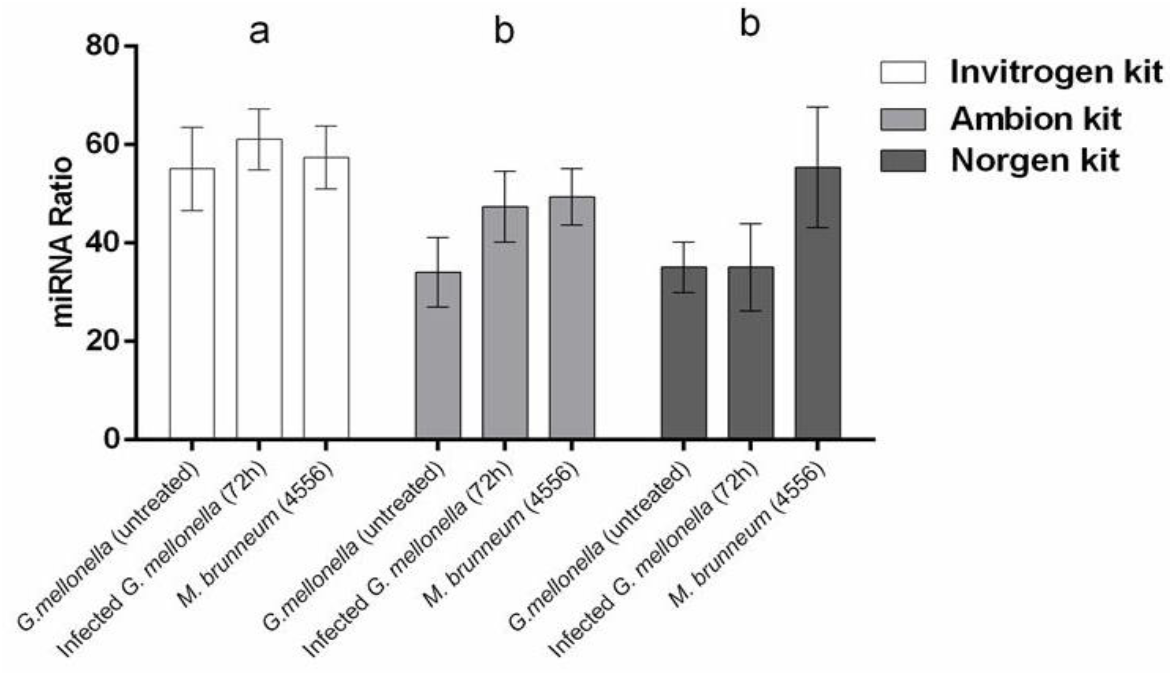
miRNA to small RNA ratio quantification from each sample obtained using the different extraction kits. Error bars indicate ±SEM, different letters above bars indicate significant differences (p<0.05, ANOVA with Tukey HSD) in the miRNA ratio between kits.

**Figure. 3.**
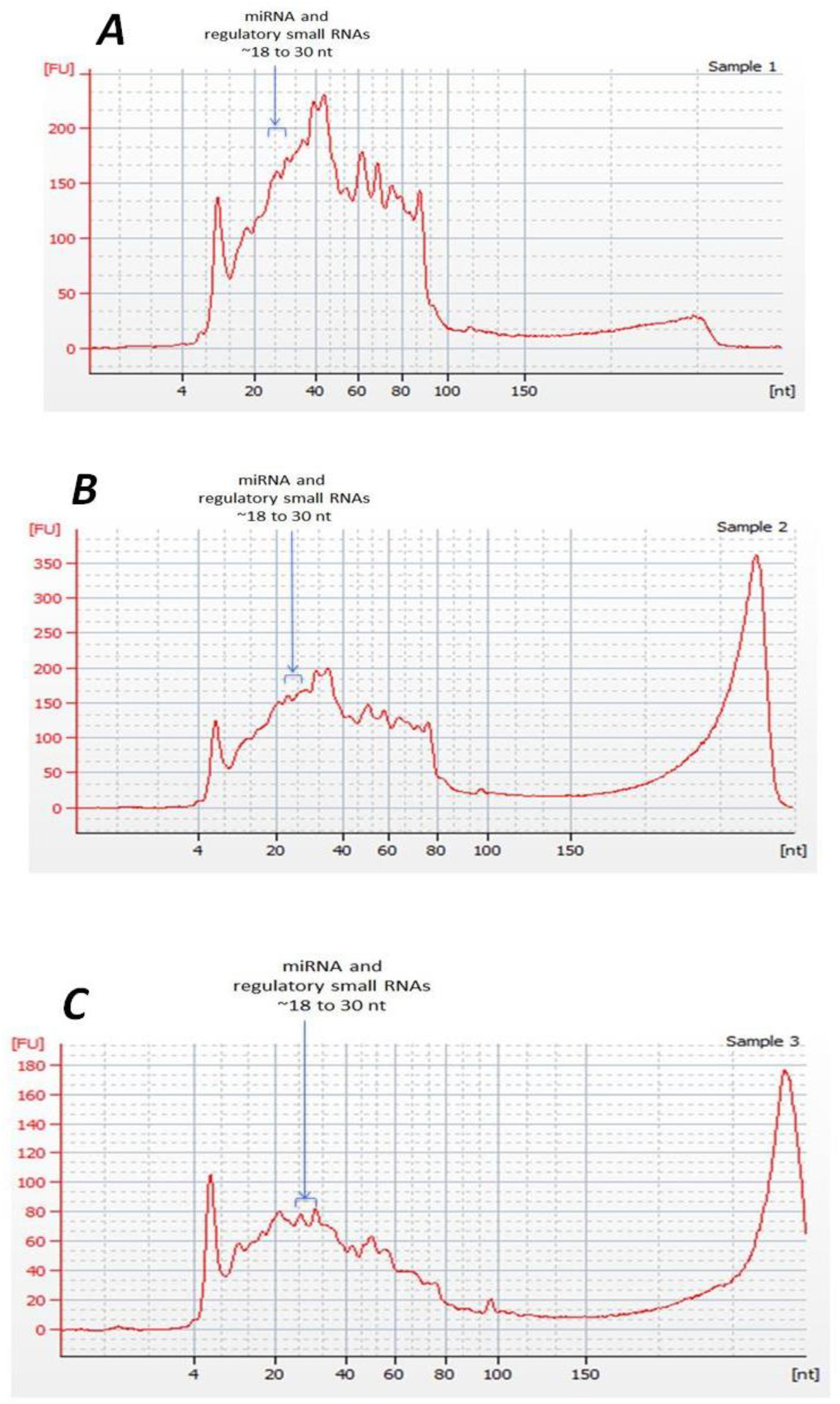
Image of a typical electropherograms for miRNAs analysis performed with the Small RNA Assay on the 2100 Bioanalyzer. Data presented from infected *G. mellonella* with *M. brunneum* for miRNA isolations using Invitrogen (A), Ambion (B) and Norgen (C) kits.

**Figure. 4.**
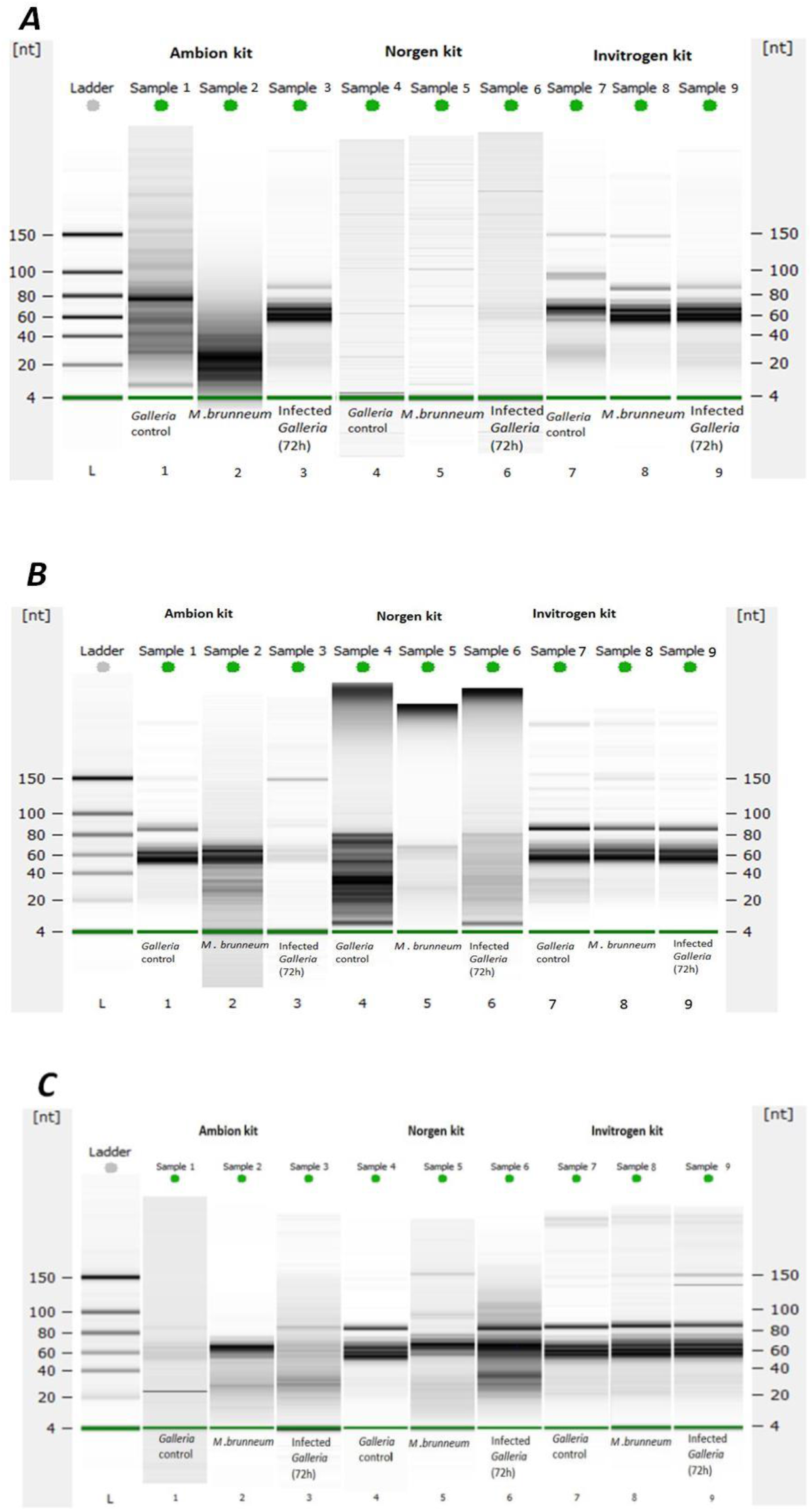
Bioanalyzer electronic gel image showing miRNA extracts using commercially available miRNA extraction kits from. A, B and C represent replicate densitometry plots for each extraction kit for the treatments *M. brunneum* (ARSEF 4556, contol), *G. mellonella* (control) and infected *G.mellonella* (72hrs).

## Discussion

High throughput molecular investigations of complex biological systems are dependant on the quality of material extracted from samples where mistakes or poor sample quailty can be expensive. Sample preparation and subsequent downstream processing and analysis have been made almost routine by proprietary kit-based protocols that offer reliability and consistency. While the rationale for the selection of a company’s kit method is not always presented by researchers, previous work on pure cultures and cell lines have shown the importance of comparing technologies when isolating microRNAs [23, 24]. The improvement in yield using a column-based protocol over non-column-based approaches (e.g. Trizol) are documented [25]. A fungal pathogen interacting within host tissues will provide specific challenges to miRNA extraction that less complex cell culture samples will not, e.g. disrupting fungal cells to obtain intact RNA, elevated presence of nucleases, diverse biochemistry. Our work emphasises the importance of correct kit selection when considering a more complex system for which high quantity, undegraded RNA is required for downstream high throughput sequencing and miRNA analysis.

While using a comparison of miRNAs from *M. brunneum*-infected *G. mellonella* larvae, *M. brunneum* culture and uninfected *G. mellonella* samples, we have shown that the selection of RNA extraction kit could have important consequences for subsequent miRNA sequencing and analysis. Such conisderations should be relevant to any plant or animal pathogen study. The kits we selected (Invitrogen PureLink, Ambion mirVana and Norgen microRNA extraction protocols) were evaluted using the protocols prescribed by the manufacturers to obtain the best results, and no modifications were made to optimize or otherwise alter the protocols. This comparison allowed us to identify a kit that not only provided the highest miRNA yield, but also had good quality miRNA and total RNA from infected *G. mellonella* larvae, consistent with non-infection controls. The Invitrogen kit was selected for our experiments because it had the highest small RNA yield and it was the easiest to use. We have shown that the Invitrogen kit produced the highest yield of microRNA (e.g. 117 ng/μl from *M. brunneum*-infected *G. mellonella* larvae) and better A_260_:A_280_ ratios (>1.9) compared to the Norgen (1.6 ng/μl) and Ambion (8.0 ng/μl) kits. While the low ratios can result from low concentration of extracted RNA [26], other studies also recorded that both Ambion and Norgen protocols yielded a similar miRNA quantity (sample extracted from pure human cell lines) in line with our findings on whole organisms and pathogen-infected cultures [4, 12].

The Invitrogen PureLink protocol combined silica column-based extraction protocol with ethanol RNA percipitation and guanidine isothiocyanate protection from degradation from RNAses. A similar process is described for the Norgen MicroRNA extraction kit, except a proprietary resin replaced silica in the column. In addition the Norgen kit is more limited in the amount of tissue that can be processed per sample (50 mg) and required more sample handling, e.g. passing the supernatant through a filter cartridge via centrifugation. The Ambion mirVana protocol is fundamentally different employing a phenol:chlorofrom extraction and ethanol preciptation and use of glass fibre-based filtration. Phenol use and disposal places an additional consideration for some laboratories. While it is not clear whether the differences in extraction chemistry resulted in the different extraction values between the kits, the lower level of RNA degradation observed for the Invitrogen PureLink kit suggests that the reduced handling time of 15 minutes per sample could be a key factor (NB samples were extracted at the same time to increase efficieny so each individual extracted sample was less than 15 or 30 mintues as recorded). Improved yield and quality may have been obtained for each extraction kit following in depth optimisation, but in conclusion our findings showed that the Invitrogen PureLink™ miRNA Isolation Kit offers more precision in extracting sequencing quality miRNA from insect and fungal tissues without the need for further optimisation.

## Conclusion

By trialing different commercially available miRNA extraction kits, we have shown variation in terms of isolated miRNA quality, quantity and reproducibility between protocols when extracting from complex tissues, namely insect larvae parasitised by a pathogenic fungus. We demonstrated that, for our experiments, the Invitrogen PureLink™ miRNA Isolation Kit provided the highest quality and quantity of miRNA to allow high throughput sequencing of the sample. Also the miRNA obtained via Ambion and Norgen kits showed a greater amount of degradation. In addition the Invitrogen protocol was technically simpler with fewer steps and did not use phenol. Therefore, while we recommend that researchers extracting miRNA from complex / environmental samples should consider testing different commercial protocols when optimising their methodology, in our hands the Invitrogen PureLink™ miRNA Isolation Kit worked well with a mixed insect-fungal pathogen system.

## Declarations

### Ethics approval and consent to participate

Not applicable

### Consent for publication

We confirm that this work is origenal and has not pulished elswhere, nor it is curently under considration for publication elsewhere. All authors have agreed to the submitted version of the manuscript. There are not any financial supports or relationships that may pose conflict of ineres.

### Availability of data and material

All data generated or analysed during the current study are includecd in this published article study

### Competing intrests

The authors declare that they have no competing interests.

### Athours’ contributions

MAA, TMB, DCE conceived and desigened the experiments.MAA Performed the experiments and performed the statistical analysis. MAA and DCE analyzed the data. Contributed reagents and materials MAA,TMB and DCE. MAA and DCE wrote the paper.

## Acknowledgments

College of Science, University of Tabuk, Tabuk KSA – provided funding for this research.

## References

1. Zhou Q, Wang Z, Zhang J, Meng H, Huang B: Genome-wide identification and profiling of microRNA-like RNAs from Metarhizium anisopliae during development. Fungal Biology 2012, 116(11):1156–1162.

2. Axtell MJ, Westholm JO, Lai EC: Vive la différence: biogenesis and evolution of microRNAs in plants and animals. Genome Biology 2011, 12(4):221.

3. Pritchard CC, Cheng HH, Tewari M: MicroRNA profiling: approaches and considerations. Nature Reviews Genetics 2012, 13(5):358–369.

4. Monleau M, Bonnel S, Gostan T, Blanchard D, Courgnaud V, Lecellier C-H: Comparison of different extraction techniques to profile microRNAs from human sera and peripheral blood mononuclear cells. BMC Genomics 2014, 15(1):395.

5. Jeffries MKS, Kiss AJ, Smith AW, Oris JT: A comparison of commercially-available automated and manual extraction kits for the isolation of total RNA from small tissue samples. BMC Biotechnology 2014, 14(1):94.

6. Wang K, Yuan Y, Cho J-H, McClarty S, Baxter D, Galas DJ: Comparing the MicroRNA spectrum between serum and plasma. PLoS One 2012, 7(7):e41561.

7. Cheng HH, Yi HS, Kim Y, Kroh EM, Chien JW, Eaton KD, Goodman MT, Tait JF, Tewari M, Pritchard CC: Plasma processing conditions substantially influence circulating microRNA biomarker levels. PLoS One 2013, 8(6):e64795.

8. Tzimagiorgis G, Michailidou EZ, Kritis A, Markopoulos AK, Kouidou S: Recovering circulating extracellular or cell-free RNA from bodily fluids. Cancer Epidemiology 2011, 35(6):580–589.

9. Kim Y-K, Yeo J, Kim B, Ha M, Kim VN: Short structured RNAs with low GC content are selectively lost during extraction from a small number of cells. Molecular Cell 2012, 46(6):893–895.

10. Arroyo JD, Chevillet JR, Kroh EM, Ruf IK, Pritchard CC, Gibson DF, Mitchell PS, Bennett CF, Pogosova-Agadjanyan EL, Stirewalt DL: Argonaute2 complexes carry a population of circulating microRNAs independent of vesicles in human plasma. Proceedings of the National Academy of Sciences 2011, 108(12):5003–5008.

11. Turchinovich A, Weiz L, Langheinz A, Burwinkel B: Characterization of extracellular circulating microRNA. Nucleic Acids Research 2011, 39(16):7223–7233.

12. Li Y, Kowdley KV: Method for microRNA isolation from clinical serum samples. Analytical Biochemistry 2012, 431(1):69–75.

13. Burgos KL, Javaherian A, Bomprezzi R, Ghaffari L, Rhodes S, Courtright A, Tembe W, Kim S, Metpally R, Van Keuren-Jensen K: Identification of extracellular miRNA in human cerebrospinal fluid by next-generation sequencing. RNA 2013, 19(5):712–722.

14. Zimmermann G: Review on safety of the entomopathogenic fungi Beauveria bassiana and Beauveria brongniartii. Biocontrol Science and Technology 2007, 17(6):553–596.

15. Ferron P: Biological control of insect pests by entomogenous fungi. Annual Review of Entomology 1978, 23(1):409–442.

16. Roush R, Tabashnik BE: Pesticide resistance in arthropods: Springer Science & Business Media; 2012.

17. Mylonakis E, Moreno R, El Khoury JB, Idnurm A, Heitman J, Calderwood SB, Ausubel FM, Diener A: Galleria mellonella as a model system to study Cryptococcus neoformans pathogenesis. Infection and Immunity 2005, 73(7):3842–3850.

18. Vilcinskas A, Götz P: Parasitic fungi and their interactions with the insect immune system. Advances in parasitology 1999, 43:267–313.

19. Kaito C, Sekimizu K: A silkworm model of pathogenic bacterial infection. Drug Discov Ther 2007, 1(2):89–93.

20. Vega FE, Goettel MS, Blackwell M, Chandler D, Jackson MA, Keller S, Koike M, Maniania NK, Monzon A, Ownley BH: Fungal entomopathogens: new insights on their ecology. Fungal Ecology 2009, 2(4):149–159.

21. Dubovskii I, Grizanova E, Chertkova E, Slepneva I, Komarov D, Vorontsova YL, Glupov V: Generation of reactive oxygen species and activity of antioxidants in hemolymph of the moth larvae Galleria mellonella (L.)(Lepidoptera: Piralidae) at development of the process of encapsulation. Journal of Evolutionary Biochemistry and Physiology 2010, 46(1):35–43.

22. Field A: Discovering statistics using IBM SPSS statistics: Sage; 2013.

23. Burgos KL, Javaherian A, Bomprezzi R, Ghaffari L, Rhodes S, Courtright A, Tembe W, Kim S, Metpally R, Van Keuren-Jensen K: Identification of extracellular miRNA in human cerebrospinal fluid by next-generation sequencing. Rna 2013, 19:712–722.

24. Li Y, Kowdley KV: Method for microRNA isolation from clinical serum samples. Analytical biochemistry 2012, 431:69–75.

25. Duy J, Koehler JW, Honko AN, Minogue TD: Optimized microRNA purification from TRIzol-treated plasma. BMC genomics 2015, 16(1):95.

26. Desjardins P, Conklin D: NanoDrop microvolume quantitation of nucleic acids. Journal of Visualized Experiments: JoVE 2010(45).

